# Specific transcriptomic signatures and dual regulation of steroidogenesis between fetal and adult mouse Leydig cells

**DOI:** 10.1101/2021.04.14.439819

**Authors:** Pauline Sararols, Isabelle Stévant, Yasmine Neirijnck, Diane Rebourcet, Annalucia Darbey, Michael K. Curley, Françoise Kühne, Emmanouil Dermitzakis, Lee B. Smith, Serge Nef

## Abstract

Leydig cells (LC) are the main testicular androgen-producing cells. In eutherian mammals, two types of LCs emerge successively during testicular development, fetal Leydig cells (FLCs) and adult Leydig cells (ALCs). Both display significant differences in androgen production and regulation. Using bulk RNA sequencing, we compared the transcriptomes of both LC populations to characterise their specific transcriptional and functional features. Despite similar transcriptomic profiles, a quarter of the genes show significant variations in expression between FLCs and ALCs. Non-transcriptional events, such as alternative splicing was also observed, including a high rate of intron retention in FLCs compared to ALCs. The use of single-cell RNA sequencing data also allowed the identification of nine FLC-specific genes and 50 ALC-specific genes. Expression of the corticotropin-releasing hormone 1 (*Crhr1*) receptor and the ACTH receptor melanocortin type 2 receptor (*Mc2r*) specifically in FLCs suggests a dual regulation of steroidogenesis. The androstenedione synthesis by FLCs is stimulated by luteinizing hormone (LH), CRH and ACTH whereas the testosterone synthesis by ALCs is dependent exclusively on LH. Overall, our study provides a useful database to explore LC development and function.

## Introduction

Leydig cells (LC) are the main steroidogenic cells of the testes. They synthesise androgens that are essential for both masculinisation of the organism and spermatogenesis. In mice, two populations of Leydig cells arise sequentially, one during embryonic development referred as the fetal Leydig cells (FLCs) and the other postnatally referred as the adult Leydig cells (ALCs) (Baker et al., 1999; Habert et al., 2001; O’Shaughnessy et al., 2002; Haider, 2004; Chen et al., 2009). The mouse FLCs appear in the interstitial compartment of the testis shortly after sex determination at embryonic day (E)12,5. The FLC population expands considerably during fetal testis development through the recruitment and differentiation of Leydig progenitor cells rather than by mitotic division of differentiated FLCs (Byskov, 1986; Kerr et al., 1988; Migrenne et al., 2001; Brennan et al., 2003; Barsoum and Yao, 2010; Ademi et al., 2020). The maximum number of FLCs is reached around birth and regresses over the first two weeks of postnatal life (Kerr et al., 1988; Shima, 2019). The ALCs appear around one week after birth and increase in number during puberty. They arise from LC progenitors located in the testicular interstitium (Davidoff et al., 2004; Barsoum et al., 2013; Shima et al., 2013; Kilcoyne et al., 2014; Ademi et al., 2020). Two recent studies showed that both fetal and adult Leydig cells derive from a common pool of progenitor cells originating from the gonadal surface epithelium and mesonephric mesenchymal cells present from fetal life (Ademi et al., 2020; Shen et al., 2020). Evidence also shows that a subset of FLCs dedifferentiate at fetal stages to serve as potential ALC stem cells (Shima et al., 2018).

The rodent FLCs and ALCs have distinct morphological and functional differences. The FLCs display a high proportion of lipid droplets, while mostly absent in the ALCs (Huhtaniemi and Pelliniemi, 1992; Shima, 2019). Unlike ALCs, the FLCs are not capable of fully synthesising testosterone on their own. They express all the enzymes necessary for androgen synthesis except HSD17B3, which converts androstenedione to testosterone. The conversion of androstenedione produced by the FLCs is achieved by the adjacent fetal Sertoli cells that express HSD17B3 (O’Shaughnessy et al., 2000; Shima et al., 2013). Another notable difference between fetal and adult LCs is their regulation by the pituitary gonadotropins. Although the luteinizing hormone (LH) receptor is expressed from E16.5 in FLCs and later in ALCs (O’Shaughnessy et al., 1998)), LH signalling is dispensable for FLCs development, but prove to be essential for ALCs development and testosterone production. Neonatal mouse mutants for LH/CG receptors display testes indistinguishable from control mice. In contrast, testes from adult mutants for LH/CG receptors are reduced in size, with fewer and hypoplastic ALCs, and show impaired testosterone production (Lei et al., 2001; Zhang et al., 2001; O’Shaughnessy and Fowler, 2011; Teerds and Huhtaniemi, 2015). FLC function is normal in the absence of endogenous circulating gonadotropins (O’Shaughnessy et al., 1998) but markedly reduced in late gestation in *T*/*ebp*/*Nkx2.1* null mice lacking a pituitary gland (Pakarinen et al., 2002). This suggests that additional hypothalamo/pituitary hormones, other than LH, may be required for FLC function and androgen production. Interestingly, two additional hormones have been reported to stimulate testosterone production in fetal testis. Adrenocorticotropic hormone (ACTH) has been reported to stimulate *in vitro* testosterone production in fetal and neonatal testes (O’Shaughnessy et al., 2003). In parallel, corticotropin-releasing hormone (CRH) has been reported to stimulate steroidogenesis by direct activation of FLCs in fetal rat and mouse testes *ex vivo* and in MA-10 mouse Leydig cells (McDowell et al., 2012), but not in primary ALCs (Huang et al., 1995; McDowell et al., 2012).

While testosterone synthesis is subjected to intensive studies, our knowledge of FLCs and ALCs origins, development, and in particular similarities and differences is still incomplete. Multiple transcriptomic studies including either mouse fetal or adult Leydig populations have been performed on whole gonads or purified cell populations in a given context (Nef et al., 2005; Beverdam and Koopman, 2006; Jameson et al., 2012; Munger et al., 2013; McClelland et al., 2015; Inoue et al., 2016; Miyabayashi et al., 2017). To date, no comprehensive comparison of FLCs and ALCs has been yet performed, and the identification of discriminant transcriptional signatures would be useful in distinguishing the two LC populations. In the present study, we employed a combination of bulk and single-cell RNA sequencing (RNA-seq and scRNA-seq) analyses to compare the transcriptome of FLCs and ALCs. While deep RNA-seq on purified Leydig cell populations allows the exploration of the transcriptomic landscape of FLCs and ALCs, including low expressed genes and alternative splicing; the high resolution of scRNA-seq allows us to bypass contamination issues inherent to cell population purification methods, and to identify specific marker genes that discriminate FLCs and ALCs amongst the other testicular cells. Our results provide a comprehensive view of the FLCs and ALCs transcriptional similarities and differences, unveiling important variations in terms of gene expression level and alternative splicing between the two populations of LCs. Furthermore, our analyses uncovered FLC- and ALC-specific markers that represent useful tools to study these two steroidogenic populations. Finally, amongst the FLCs specific markers we found the corticotropin-releasing hormone receptor 1 (*Crhr1*), suggesting FLCs androgen synthesis is influenced by CRH.

## Material & Methods

### Mouse strains

Embryos were collected from timed pregnant female CD-1 outbred mice (Charles River) and heterozygous *Tg*(*Nr5a1-GFP*) transgenic male mice (Stallings, 2002). The mating plug observed the next morning is designated as E0.5 were used in this study. Animals were housed and cared according to the ethical guidelines of the Direction Générale de la Santé of the Canton de Genève (GE/57/18 30080).

### Fetal and adult Leydig cell purification

The purification of fetal Leydig cells was carried out in seven independent experiments based on a previously described experimental protocol (Nef et al., 2005; Pitetti et al., 2013; Stévant et al., 2018). The resulting cells were pooled to achieve the amount of RNA required for the preparation of the RNA sequence library (**Supplementary Table S6**). In short, adult CD-1 females were time-mated with heterozygous *Tg*(*Nr5a1-GFP*) transgenic male and checked for the presence of vaginal plugs the next morning (E0.5). On the relevant days of gestation (E18.5), females were sacrificed by CO2 inhalation and the embryos collected in PBS. The sex and the presence of the *Nr5a1-GFP* transgene in the embryos were assessed under a fluorescent binocular microscope. Testes were isolated and incubated 20 minutes with trypsin-EDTA 0.05%, mechanically dissociated with gentle pipetting, and filtered through a 70μm cell strainer to obtain single cell suspension.

Adult Leydig cells purification has been performed in four independent experiments (**Supplementary Table S6**). Hundred-day-old (P100) heterozygous *Tg*(*Nr5a1-GFP*) transgenic male mice were used for this experiment. Mice were sacrificed with Esconarkon injection and ~1 mL of blood was collected by intracardiac puncture for serum extraction. Tunica albuginea of the testes were delicately removed and testes were incubated in DMEM supplemented with collagenase (1 mg/mL C0130; Sigma-Aldrich, St. Louis, MO), hyaluronidase (2 mg/mL H3506; Sigma-Aldrich), and DnaseI (0.8 mg/mL dN25; Sigma-Aldrich) at 37°C for 20 minutes with gentle agitation. After two rounds of seminiferous tubules sedimentation, the supernatants enriched in interstitial cells were collected and incubated 10 minutes with Trypsine-EDTA 0.05%. Cells were centrifuged and filtered through a 70μm cell strainer to obtain single cell suspension.

Nr5a1-GFP^+^ cells from E18.5 and P100 testes were then sorted by fluorescent-active cell sorting (BD FACS ARIA II), excluding cell doublets, and the dead cells with Draq7TM dye staining. Cells were collected directly into RLT buffer from Qiagen RNeasy Mini kit for RNA extraction.

### Bulk RNA-sequencing library preparation and sequencing

RNA was extracted from Nr5a1-GFP^+^ cells with the RNeasy Mini kit (Qiagen) to obtain a minimum of 260 ng of total RNA. The composition of the different samples is detailed in **Supplementary Table S6**. Sequencing libraries were prepared from 150 ng of DNA with the TruSeq Stranded Total RNA Library Prep Gold (Ribo-Zero) and sequenced on an Illumina HiSeq 2500 (50 bp, paired end, ~35 million reads expected) at the Genomics Platform of the University of Geneva.

### RNA-sequencing analysis

Reads were demultiplexed with Casava (v1.8.2), mapped with GemTools (v1.7.1) (Marco-Sola et al., 2012) and read counts and RPKM gene expression quantifications were calculated with an in-house pipeline based on Gencode annotation GRCm38 (v4). Globally, over 88% of the reads mapped to exonic regions. Pre-ranked gene set enrichment analysis was performed using GSEA (v4.1.0) with the mean RPKM expression as ranking, and using the genes with a mean RPKM>1 (Subramanian et al., 2005). Spearman correlation and principal component analysis (PCA) using the R base stats package, revealed a very high correlation for both biological triplicates of ALCs and FLCs (Spearman correlation score >98%, see **Supplementary Figure 1C** and **D**). Similarly, the correlation between the two conditions, ALCs and FLCs, is also very high (Spearman correlation score >88%). We used the R package DESeq2 (v1.24.0) (Love et al., 2014) for the differential expression analysis. Genes with fewer than 10 reads were not taken into account. GO terms enrichment analysis was computed on the selected genes enriched in FLCs and in ALCs with the R packages ClusterProfiler (v3.12.0) (Yu et al., 2012).

Differential splicing analysis was performed using rMATS (v3.2.5) (Shen et al., 2012, 2014) with the bam files as input. Splicing event with an FDR<=0.05 were considered as significant.

### Single-cell RNA-sequencing library preparation and sequencing

Single-cell library of E16.5 testis from Neirijnck et al. (Neirijnck et al., 2019) (**GSE123119**) was prepared with the Chromium Single Cell 3’ Library v2 kit and sequenced targeting 5,000 cells with an Illumina Hiseq 4000 (100 bp, paired-end, 100,000 reads per cell expected) with Macrogen (http://foreign.macrogen.com/eng/).

Adult testis from adult mice (C57BL/6J mice) single-cell RNA sequencing data come from Ernst et al. (Ernst et al., 2019) (**E-MTAB-6946**). We used the libraries do15983, do15984, do17622, do17623, do17815, do17816, do18197, do18198, do18199 of mice older than 60 days old from Ernst et al. The libraries were prepared with the Single Cell 3’ Library v2 kit and sequenced with an Illumina HiSeq 2500.

### Data processing with the Cell Ranger package, cell selection and in-house quality controls

Computations were performed at the Vital-IT Centre for high-performance computing of the SIB (Swiss Institute of Bioinformatics) (http://www.vital-it.ch). Demultiplexing, alignment, barcode filtering and UMI counting were performed with the Cell Ranger v2.1 pipeline (10x Genomics). Reference genome has been modified to include the eGFP transgene with the mkref function. Data were mapped to the mouse reference genome GRCm38.p5 in which the *eGFP* and the bovine GH 3’ splice/polyadenylation signals (NM_180996.1) (Stallings, 2002) sequences have been added, and annotated with Gencode vM15. Only protein coding genes and long non-coding RNAs were retained for further analysis.

To set a threshold between cell containing barcodes and empty ones, we computed the knee point and the inflection point of the ranked barcode distribution plot. Then, we detected the local minimum between these points on the density curve (density base R function, bw=500, n=4,096) of the UMI number per barcode using the quantmod R package (v0.4-16). This local minimum was used as a threshold for cell selection.

### Single-cell RNA-sequencing analysis

#### E16.5 mouse RNA-sequencing data

The single-cell RNA sequencing analysis of the E16.5 testis (Neirijnck et al., 2019) were performed using the Seurat software package (v2.3.4). From the raw matrix obtained with Cell Ranger version 2.0 (10X Genomics), we filtered cells based on the UMI count per cell (>2300 UMI) and on the percentage of mitochondrial genes (>0.05% of mitochondrial genes) resulting in 3,781 cells. Then, we reduced the size of the dataset using Principal Component Analysis (PCA) on the genes expressed in more than 50 cells (9,576 genes) and calculated the UMAP representation (Becht et al., 2019) and finally grouped the cells with Louvain algorithm (Waltman and van Eck, 2013) (resolution=1) using the 15th first PCs. We classified the clusters according to the marker genes of the different cell types in the literature, including the genes *Nr5a1*, *Star*, *Cyp11a1*, *Insl3*, *Hsd17b3*, *Amh*, *Sox9*, *Arx*, *Nr2f2*, *Pecam1*, *Cdh5*, *Esam*, *Pou5f1*, *Ddx4* (Castrillon et al., 2000; Wakayama et al., 2003; Okamura et al., 2008; Yu et al., 2009; Buaas et al., 2012; Wen et al., 2016; Kumar et al., 2017; Stévant et al., 2018; Ernst et al., 2019; Mucenski et al., 2019) and identified cluster 7 as Leydig cells.

#### Adult mouse single-Cell RNA-Sequencing data

We selected the libraries do15983, do15984, do17622, do17623, do17815, do17816, do18197, do18198, do18199 of mice older than 60 days old from Ernst et al. (Ernst et al., 2019). These libraries were analysed with Cell Ranger v1.3.1 software using the default threshold to obtain high-quality cells with large numbers of UMIs. We filtered out cells with <500 UMI and we excluded cells with more than 5% of reads mapping to the mitochondrial genome. We selected only protein-coding genes. Then, we inferred cell labels with the annotation furnished by Ernst et al., and removed cells labelled as “Outliers”. So, 24,672 cells were used. We confirmed the cell type classification of Ernst et al. with genes known from the literature (*Cyp11a1*, *Star*, *Insl3*, *Cyp17a1*, *and Fabp3*) (Stévant et al., 2018; Ernst et al., 2019).

#### Merge of Fetal and Adult single-cell RNA Sequencing data

In order to compare the Leydig cells present at fetal and adult stages, we merged the single-cell RNA sequencing datasets with the MergeSeurat function (Seurat, v2.3.4). We normalised and computed the Principal Component Analysis (PCA) using the genes expressed in more than 50 cells. The 10 first components of this PCA were used to compute the corrected neighbour graph with BBKNN (balanced batch KNN, v1.3.8) (Polański et al., 2019) and then, the umap representation. We made use of the previous cell annotation to distinguish fetal and adult populations and we ensured the cell identity using marker gene expression. This list includes the genes *Nr5a1*, *Star*, *Cyp11a1*, *Insl3*, *Hsd17b3*, *Amh*, *Sox9*, *Arx*, *Nr2f2*, *Pecam1*, *Cdh5*, *Esam*, *Pou5f1*, *Ddx4*, *Dmrt1*, *Piwil1*, *Pex21*, and *Tnp1*. We performed a differential expression analysis (Mann-Whitney Wilcoxon test) between FLCs and ALCs clusters using Seurat FindMarkers function (only positive markers, min.pct=0.25, thresh.use=0.25). We identified genes showing a high expression in the one population of Leydig cells and a low expression in the other population (adj. p-valuej<0.01, avg logFC>0.5, pct.1>0.5, pct.2<0.25), and overexpressed as well in the same population in the DESeq2 analysis.

From this selection, in order to select the genes with specific expression in one Leydig cell population and with a low expression in all other populations in the testis, we used an additional differential expression analysis between all cell types using Seurat FindAllMarkers function (only positive markers, p_val_adj<0.01 & avg_logFC>0.5 & pct.1>0.5 & pct.2<0.25) was used to compute marker genes for every cluster. The intersect of the two differential expression analysis with Seurat is used to get a list of the marker genes of FLCs and ALCs specifically.

### RNAscope^®^ analyses (*In situ* hybridization)

Adult (P100) and embryonic (E16.5) *Nr5a1-eGFP* samples from timed mated females were collected and fixed overnight in 4% paraformaldehyde, dehydrated and embedded in paraffin. Five μm thick sections were examined histologically *via* haematoxylin and eosin staining. We performed the RNAscope^®®^ 2.5 HD DuplexAssay protocol following the recommendation of BioTechne. The *Star* probes (C2) to label Leydig cells, and the probes for the candidates *Crhr1* (C1), *Ren1* (C1), *Bhmt* (C1) and *Sult1e1* (C1) were tested. Slides were imaged using an Axioskop 2 plus confocal microscope and ZEN 2009 software (Carl Zeiss Ltd, Hertfordshire, UK). For reproducibility purpose, at least three different animals of each group were tested.

### Immunostaining

Animals were bred and maintained in strict compliance with the Animals (Scientific Procedures) Act, 1986. All procedures were conducted in accordance with United Kingdom Home Office regulations under project licenses 60/4200 and 70/8804 held by Lee B. Smith.

Neonatal and adult tissues were fixed in Bouins for 6 hours, stored in ethanol 70% and embed in paraffin. Sections of 5 μm were dewaxed in xylene, rehydrated in graded ethanol solutions. For the double immunostaining, slides were antigen-retrieved in pressure cook with 0.01M citrate buffer (pH 6.0). To quench endogenous peroxidases activity, slides were incubated in 0.3% hydrogen peroxide (v/v) in TBS for 30 min at room temperature (RT). The non-specific activity was blocked using the appropriate normal blocking serum for 30 min at RT followed by incubation overnight at 4°C with the first primary antibody diluted in blocking serum. After washing, slides were incubated for 30 min at RT with the appropriate secondary antibody conjugated to peroxidase and diluted 1/200 in blocking serum and left on the slides for 30 min at RT. Sections were then incubated with Tyramide Signal Amplification system (‘TSA^™^’, Perkin Elmer) diluted 1/50 for 10 min at RT according to the manufacturer’s instructions. Slides were then stained with the second Primary antibody and washed as an incubated as described above with secondary and Tyramide. Sections were then counterstained in Sytox Green (Molecular Probes, life technologies, Paisley, UK) for 10 min at RT and mounted in PermaFluor mounting medium (Thermo Scientific, UK). Slides were scanned using an LSM 710 confocal microscope and ZEN 2009 software (Carl Zeiss Ltd, Hertfordshire, UK). The primary and adequate secondary antibodies used in this study are detailed in **Supplementary Table S7**. To assure the specificity of the stained tissue, sections incubated with no primary antibody were used as negative controls. For reproducibility purpose, at least three different animals of each group were tested.

### Data Availability

The fetal and adult Leydig cell bulk RNA-seq data are available on GEO (NCBI) under accession number **GSE171746**.

## Results

### A global view of fetal and adult Leydig cell transcriptomes

To compare the transcriptomic signatures of FLCs and ALCs, we purified Leydig cells at embryonic day (E) 18.5 and postnatal day (P) P100. To proceed, we FAC-sorted the highest GFP positive cell population corresponding to Leydig cells from *Nr5a1*-*eGFP* testes (Stallings, 2002) (**Figure 1A; **Supplementary Figure 1A** and A’**). We then performed RNA sequencing of poly(A)^+^ RNAs in biological triplicates for the two stages. In total, we identified 20,859 and 21,195 expressed genes in FLCs and ALCs respectively (with RPKM≥1), with 78.5% (±0.5) of them being protein coding. We evaluated the purity of our FLCs and ALCs samples by measuring the expression level of several genes specific to different testicular cells (**Supplementary Figure 1B**). Our samples display high expression levels of the Leydig cell marker genes *Cyp11a1*, *Insl3*, *Cyp17a1*, *Hsd3b1*, *and Star* (Rebourcet et al., 2019). In contrast, expression of marker genes for Sertoli cells (*Sox9*, *Dhh*, *Amh*) (Bitgood et al., 1996; Liu et al., 2016; Rehman et al., 2017), interstitial progenitors cells (*Nr2f2*, *Arx*, *Tcf21*), germ cells (*Dazl*, *Ddx4*, *Pou5f1*) (Castrillon et al., 2000; Okamura et al., 2008; Yu et al., 2009), and endothelial cells (*Tek*, *Pecam1*, *Esam*) (De Val and Black, 2009; Stévant et al., 2018), immune cells (*Cxcl2*, *Ptprc*, *Coro1a*), and peritubular myoid cells (*Acta2*, *Myh11*, *Cnn1*) were low, confirming the high degree of enrichment of our FLC and ALC samples (Chen et al., 2014; Rebourcet et al., 2014).

**Figure 1:**
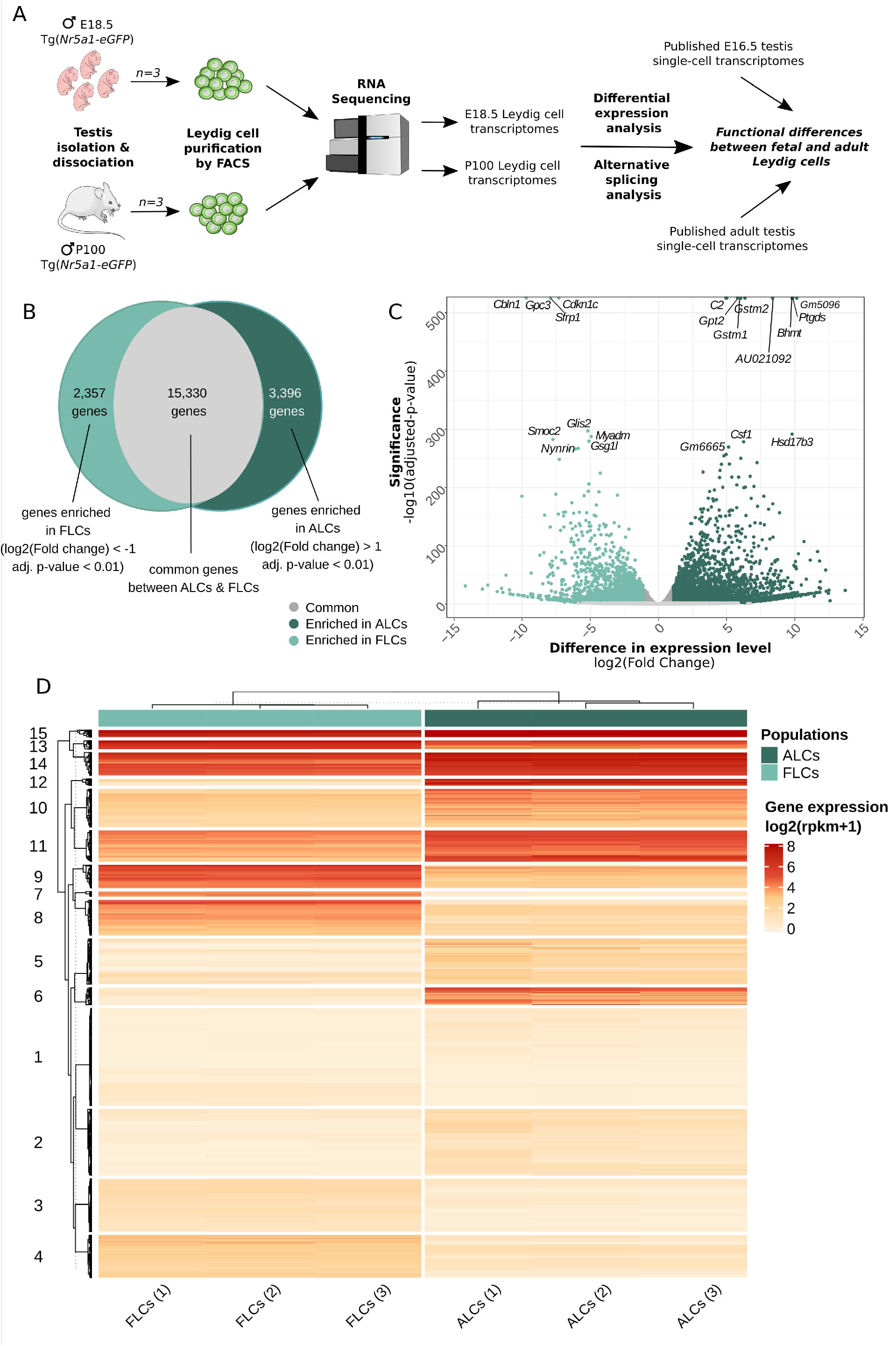
Similarities and differences between FLCs and ALCs. **(A)** Sample collection, RNA-sequencing and analysis workflow. **(B)** Venn diagram showing the number of genes expressed at same levels in FLCs and ALCs (12,645 genes), enriched in FLCs (3,741 genes) and enriched in ALCs (4,657 genes). **(C)** Volcano plot displaying differential expressed genes between FLCs and ALCs. The x-axis corresponds to the log2(Fold Change) and the y-axis corresponds to the −log10(adjusted p-value). The salmon dots represent the significant up-regulated transcripts in FLCs (adj. pval<0.01); the blue dots represent the significant up-regulated transcripts in ALCs (adj. pval<0.01) and the grey dots represent the the not significantly differentially expressed transcripts (adj. pval>0.01). Top 30 genes according to the adjusted p-value are displaying on this volcano plot. **(D)** Heatmap displaying the normalized scaled expression of differentially expressed genes in the 6 samples. Genes are ordered with hierarchical clustering according to their expression pattern into 15 groups.

We show that the ALCs express in excess the *Insl3* gene with an average of 21,113 RPKM, followed by *Aldh1a1* (8,050 RPKM) and *Cyp17a1* (5,742 RPKM). Together the RNA abundance of three genes represents 9.4% of the whole transcriptome. The FLCs do not display such an extreme over-representation of the same genes, but the top three expressed genes are *mt-Co1* (7,501 RPKM), *Hsd3b1* (5,357 RPKM), and *Insl3* (3,947 RPKM), which represent 5.4% of the total transcriptome. To appreciate the biological functions enriched amongst the most expressed genes of FLCs and ALCs, we performed a pre-ranked gene set enrichment analysis of the transcriptomes, weighting genes by their level of expression. While in FLCs we observed a statistical enrichment of 345 GO terms (FDR<25%), including “glucocorticoid biosynthetic process”, “C21 steroid hormone metabolic process” and “regulation of systemic arterial blood pressure by renin angiotensin” in the top terms, no statistical enrichment was observed for the ALCs. Although not statistically enriched, we could find in the top terms “glucocorticoid metabolic process”, “C21 steroid hormone metabolic process”, and “circadian sleep wake cycle” (of note, the P100 mice were euthanized in the morning, and we know testosterone synthesis is sensitive to the circadian rhythm (Chen et al., 2017)) (**Supplementary Table S2**). These results indicate that both fetal and adult Leydig cells are highly specialised cells dedicated to steroid production. For all subsequent analyses, we retained only the genes coding for proteins and long non-coding RNA (lncRNA).

### Wide variations in gene expression levels were observed between FLCs and ALCs

We then thought to evaluate the extent to which the transcriptomes of the two Leydig cell populations are comparable and what genes and biological pathways are differentially expressed. Among the 21,083 protein coding and long-non-coding genes expressed in either FLCs or ALCs, 15,330 genes have no significant difference of expression (**Figure 1B** and **Supplementary Table S1**). Interestingly, a large proportion of genes (5,753 genes, *i.e*. 27% of the total) display significant variations of expression levels between ALCs and FLCs (adjusted p-value<0.01). Of these genes, 2,357 are overexpressed in FLCs (FC>2) (**Figure 1B** and **C**). The vast majority of genes overexpressed in FLCs have never been identified as such. This is particularly the case for the top 30 genes with the lowest adjusted p-value such as *Cdkn1c*, *Gpc3*, *Cbln1*, *Sfrp1*, *Myadm*, *Glis2*, *Peg3* and *Smoc2*. In addition, we found also *Thbs2 and Mc2r*, two genes known to be specifically expressed in FLCs (O’Shaughnessy et al., 2002, 2003) as well as 50 genes already reported to be expressed - although not specifically - in FLCs (Jameson et al., 2012; McDowell et al., 2012; McClelland et al., 2015; Inoue et al., 2016)). We grouped the genes differentially expressed into 15 clusters according to their expression profile using a hierarchical clustering (**Figure 1D**). Genes enriched in FLCs are grouped in clusters 3, 4, 7, 8, 9, and 13. Gene Ontology (GO) analysis of these clusters revealed an association with the development of the urogenital system (cluster 3), cell division and differentiation (cluster 4), various metabolic processes (clusters 7, 9, 13), and response to reactive oxygen species (ROS) (cluster 13) (**Supplementary Table S2**). On the other side, 3,396 genes were found overexpressed in ALCs (FC<0.5) (**Figure 1B** and **C**). Again, the large majority of genes overexpressed in ALCs have never been identified as such. This is particularly the case for genes with the lowest adjusted p-value such as *Gstm2*, *Gstm1*, *Amy1*, *Csf1*, *Timp2*. As expected, we have found genes known to be specifically expressed in ALCs such as *Hsd3b6*, *Hsd17b3*, *Vcam1*, *Sult1e1* and *Hpgds* (O’Shaughnessy et al., 2002), as well as 750 genes already reported to be expressed - although not specifically - in ALCs (Sanz et al., 2013; O’Shaughnessy et al., 2014). Genes enriched in ALCs are grouped in clusters 2, 5, 6, 10, 11, 12, 14 and 15. This time, GO analysis of these clusters indicated a link with fertilisation (clusters 2, 5), regulation of cellular response (cluster 10), cell-substrate adhesion (cluster 11), regulation of ROS (cluster 14), and various metabolic processes (clusters 12, 14, 15) (**Supplementary Table S2**). The cluster 1 regroups genes with low expression in both LC populations that are involved in stress and immune response. This association with the immune system is consistent with the role of cytokines secreted by testicular macrophages in the regulation of Leydig cell functions (Hales, 2002). Overall, we observed significant variations in the level of expression of thousands of genes, with only a handful of genes exhibiting specific expression in one of the two Leydig cell populations.

### Differential splicing between FLCs and ALCs

Alternative splicingis a ubiquitous regulatory mechanism that allows the generation of multiple transcript isoforms from a single gene, thus expanding the complexity of the proteome. However, the extent of alternative splicing occurring in FLCs and ALCs and its functional relevance remain unclear. To investigate whether these two cell populations exhibit different alternative splicing profiles, we performed a multivariate analysis of transcript splicing. We found 1,971 splicing events that are statistically different between the two LC populations (FDR<0.05) (1,380 events in FLCs and 591 in ALCs) (**Figure 2A, Supplementary Table S2**). These splicing events occur in 1,437 genes, including 1,036 genes in FLCs, and 509 genes in ALCs (with 31 genes having an alternative splicing in both cell populations). We also examined if the genes detected as alternatively spliced correspond to differentially expressed genes. Of the 2,357 FLCs overexpressed genes, 86 of them show an alternative splicing. In the ALCs, 56 genes out of the 3,396 overexpressed genes show an alternative splicing. It appeared that the genes involved in steroidogenesis are not subject to alternative splicing in both FLCs and ALCs.

**Figure 2:**
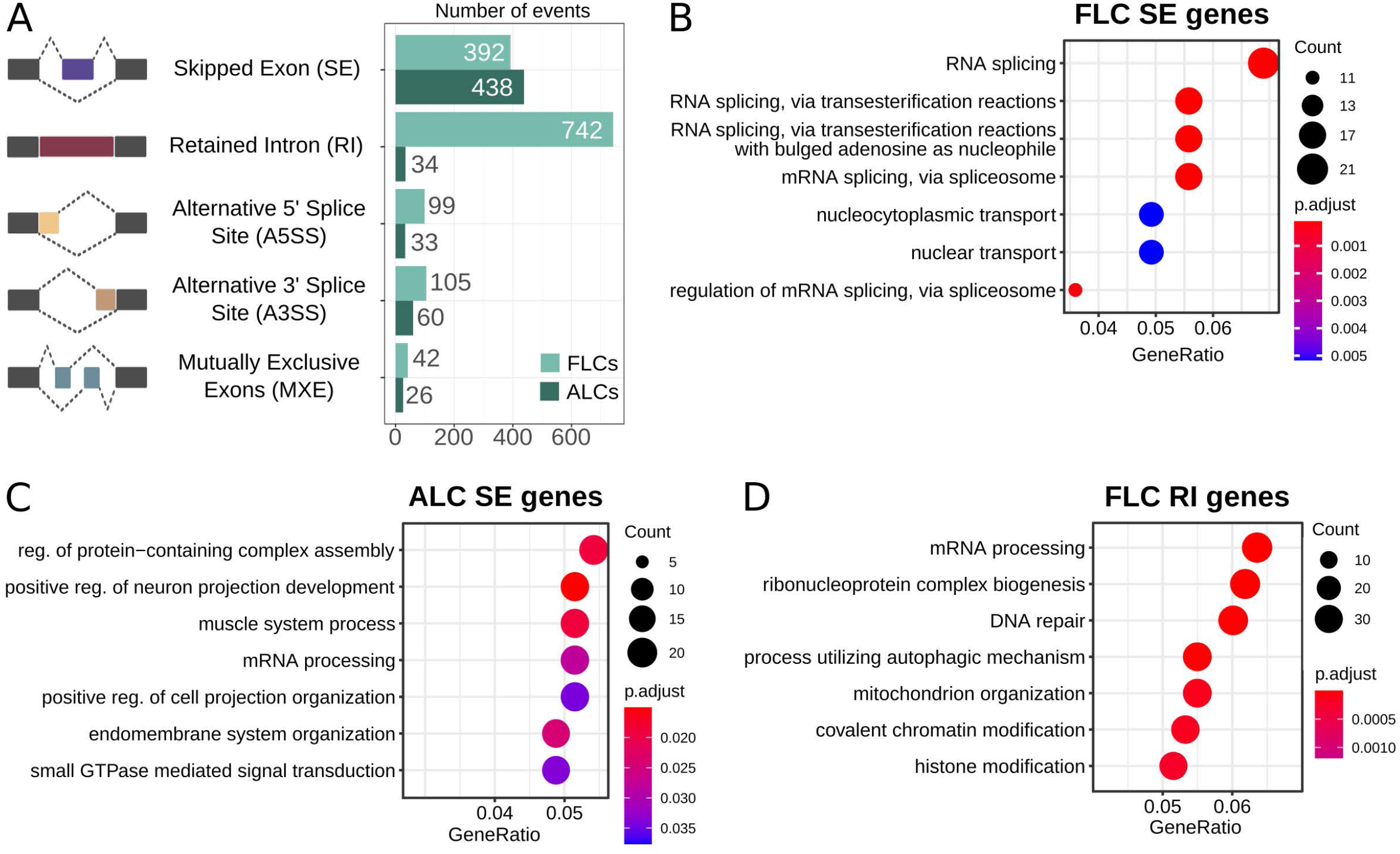
Alternative splicing. **(A)** Proportion of the differential alternative splicing events found in fetal and adult Leydig cells. **(B**, **C** & **D)** GO term enrichment of the biological functions in genes showing differential skipping exons in fetal and adult Leydig cells **(B, C)**, and intron retention in fetal Leydig cells (**D**).

As shown in **Fig. 2A**, intron retention is the most represented type of alternative splicing in the FLCs, with 742 events found in FLCs but only 34 in ALCs. We investigated if the genes presenting the different type of alternative splicing in both populations are enriched in a particular biological function by performing a GO enrichment analysis (**Supplementary Table S3**). In FLCs, the genes presenting exon skipping are strongly enriched in RNA splicing functions (**Figure 2B**), while genes showing intron retention are involved in mRNA processing and chromatin rearrangements (**Figure 2C**). Regarding ALCs, genes showing exon skipping are involved in various processes such as cellular organization, cellular projection, or muscle system process (**Figure 2D**). No GO enrichment was found in the other types of alternative splicing due to the small number of genes. Overall, we showed that intron retention is a landmark of the FLC transcriptome. It is known that alternative splicing is frequent during embryonic development, usually cell/organ specific and plays a role in gene expression regulation and protein diversity (Revil et al., 2010; Kalsotra and Cooper, 2011; Grabski et al., 2021).

### Characterisation of mutually exclusive marker genes/signatures of FLCs and ALCs

A thorough identification of the genes specifically expressed in FLCs and ALCs has never been achieved. Although our comparative analysis identified many genes that are differentially expressed between FLCs and ALCs, there is no evidence that these are specific to FLCs or ALCs. Indeed, many of them may also be expressed in other testicular cell types. To identify FLC- and ALC-marker genes, we took advantage of existing single-cell RNA sequencing data from E16.5 testes (3,781 cells) (Neirijnck et al., 2019) and adult testes (24,672 cells) (Ernst et al., 2019). We combined these fetal and adult datasets to evaluate the gene expression specificity. Using five well-established Leydig cell markers, namely *Hsd3b1*, *Star*, *Cyp11a1*, *Cyp17a1* and *Fabp3*, we identified 151 FLCs and 148 ALCs (**Figure 3)** (Ernst et al., 2019; Rebourcet et al., 2019). By comparing the sets of genes specifically enriched in ALCs and FLCs obtained with the single-cell RNA sequencing and the bulk RNA sequencing approaches described above, we generated a high-confidence selection of 62 genes enriched in FLCs showing a link with cell proliferation and differentiation, and hormone secretion (**Supplementary Table S3**) and 120 genes enriched in ALCs related to protein processing, spermatid development, and metabolic processes (**Supplementary Table S4**).

**Figure 3:**
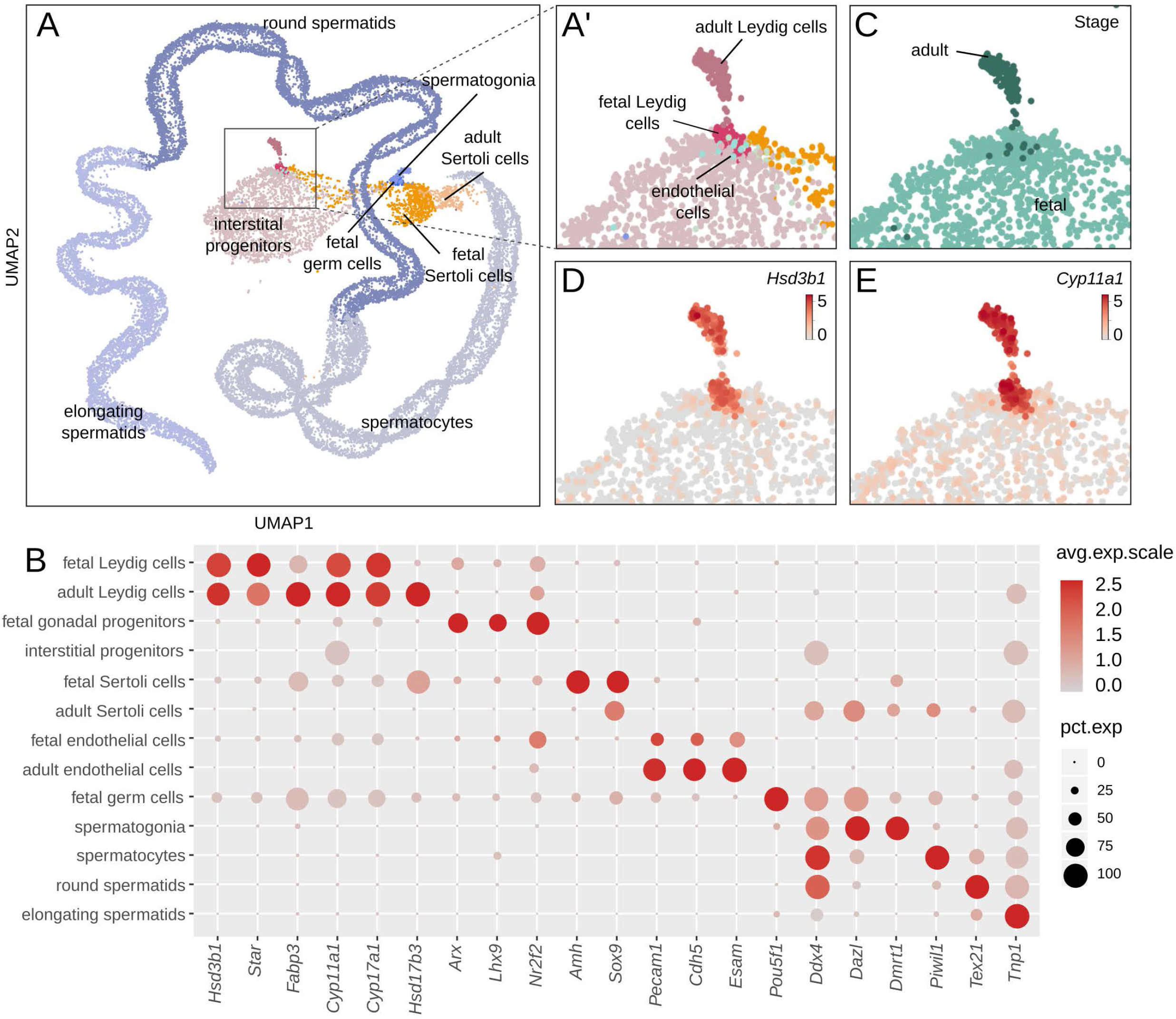
Classification of cell populations in single-cell RNA sequencing data of fetal and adult mouse testis. **(A)** UMAP representation of single-cell transcriptomes colored by cell type, where each dot corresponds to one cell. Enlargement of the global UMAP on the area of interest which include both the fetal and adult Leydig, colored by cell populations **(A’)** and by developmental stage **(C)**. The fetal cells are colored in pale green and the adult cells in dark green. **(D, E)** Enlargement on the UMAP representation colored according to the normalized expression of Leydig cells marker genes *Hsd3b1* **(D)** and *Cyp11a1* **(E)**. **(B)** Dotplot displaying the expression of selected marker genes of testis cell populations. The size of the dots is proportional to the fraction of cells in the population expressing the gene and the scaled gene expression level is indicated by the color scale. Leydig cells: *Nr5a1*, *Star*, *Cyp11a1*. Sertoli cells: *Hsd17b3*, *Amhg*, *Sox9*. Interstitial progenitors: *Arx*, *Lhx9*, *Nr2f2*. Endothelial cells: *Pecam1*, *Cdh5*, *Esam*. Germ cells: *Pou5f1*, *Dddx4*. Spermatogonia: *Dmrt1*. Spermatocytes: *Piwil1*. Round spermatids: *Tex21*. Elongating spermatids: *Tnp1*.

### Identification of nine FLC-specific marker genes

Although we have identified 62 genes showing an expression in FLCs and low or no expression in ALCs, there is no evidence that these genes are specific to FLCs. Using single-cell transcriptomic data, we excluded the ones that were also expressed in other testicular cell populations. We have thus identified nine genes that were considered as specific testicular markers of FLCs (**Figure 4A-G** and **Table 1**). This includes genes *Crhr1* (McDowell et al., 2012; McClelland et al., 2015; Inoue et al., 2016) (**Figure 4C**), *Ren1* (Jameson et al., 2012; Inoue et al., 2016) (**Figure 4D**) and *Vsnl1* (Jameson et al., 2012; McClelland et al., 2015) (**Figure 4E**), whose expression in FLCs (but not their specificity) has already been demonstrated. In addition, we have identified six additional genes described for the first time as FLC specific markers, including *Cyp26b1* (**Figure 4F**), *Gsg1l*, *Pcsk6*, *Nppc*, *Cdon* and *Ppp2r5b*. Contrary to our expectations, *Mc2r* and *Thbs2* are not part of our selection. *Thbs2* is excluded by our filters because it does not show any enrichment in FLCs compared to ALCs, while *Mc2r* (**Figure 4G**) seems specific to FLCs but is not retained due to its low expression in single-cell transcriptomic data. The specific expression of FLC marker genes *Crhr1* and *Ren1* was validated by *in situ* hybridisation and compared with *Star* expression, a known marker of Leydig cells (**Figure 4H-K**). We found that both *Crhr1* and *Ren1* are co-expressed with *Star* in E18.5 testis but not in adult P100 testis, confirming that these genes are specifically expressed in FLCs and not in ALCs or any other testicular cells (**Figure 4H-K**).

**Figure 4:**
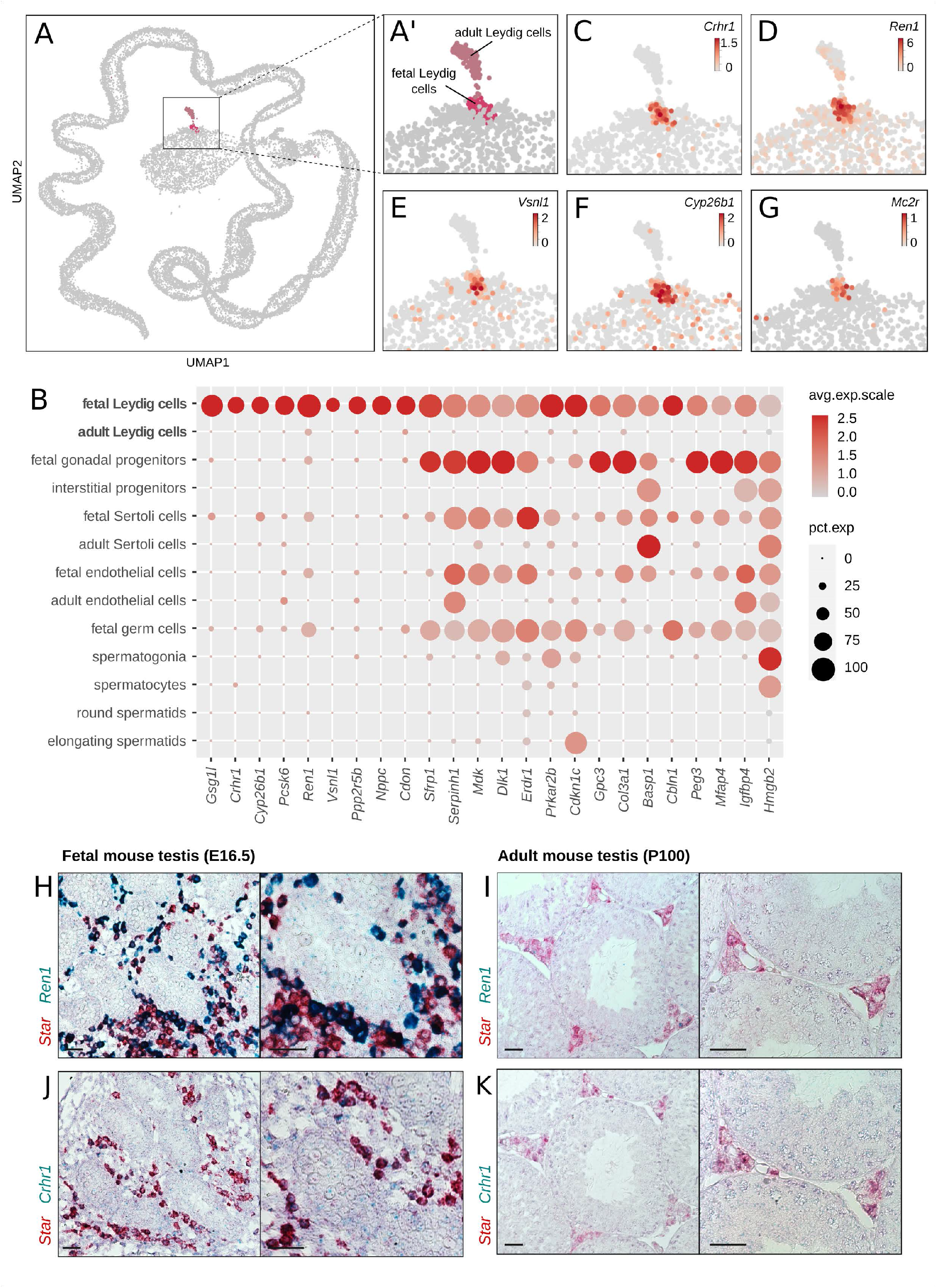
Identification of genes labelling FLCs. **(A, A’)** UMAP representation of single-cell transcriptomes highligting the two Leydig cell populations: fetal Leydig cells (FLCs) in pink and adult Leydig cells (ALCs) in grey pink. The other populations of the testis are colored in grey. **(C-G)** Enlargement on the UMAP representation colored according to the normalized expression of selected FLC-specific candidate marker genes, like *Crhr1* **(C)**, *Ren1* **(D)**, *Vsnl1* **(E)**, *Cyp26b1* **(F)**, and *Mc2r* **(G)**. **(B)** Dotplot displaying the scaled expression of the FLC-specfic candidate genes, and the top 15 non-specific candidates discriminating FLCs over ALCs. **(H-K)** *In situ* hybridization assay by RNAscope^®^ labelling in red *Star*, a known marker of Leydig cells, and in green the FLC marker *Ren1* **(H, I)** and *Crhr1* **(J, K)** in embryonic mouse testis at E18.5 **(H, J)** and in adult mouse testis at P100 **(I, K)**. The colocalisation of the red and green probes indicate a co-expression of the two genes in the Leydig cells. The black scale bar corresponds to 100μm.

**Table 1.** FLC specific genes

| Gene | baseMean | log2FoldChange | Fold Change | p value | p adj |
| --- | --- | --- | --- | --- | --- |
| *Gsg1l* | 6 453 | 5,109 | 34,511 | 2,69E-295 | 2,54E-292 |
| *Crhr1* | 3 887 | 10,022 | 1 039,829 | 4,24E-191 | 1,31E-188 |
| *Cyp26b1* | 23 010 | 6,389 | 83,795 | 1,96E-109 | 1,70E-107 |
| *Pcsk6* | 2 697 | 4,599 | 24,240 | 1,38E-56 | 3,84E-55 |
| *Ren1* | 24 384 | 5,091 | 34,091 | 1,11E-29 | 1,31E-28 |
| *Vsnl1* | 168 | 10,669 | 1 627,844 | 1,54E-17 | 1,02E-16 |
| *Ppp2r5b* | 8 531 | 1,116 | 2,168 | 1,18E-15 | 6,91E-15 |
| *Nppc* | 307 | 3,352 | 10,212 | 2,17E-14 | 1,17E-13 |
| *Cdon* | 28 637 | 2,341 | 5,067 | 1,88E-05 | 4,58E-05 |

### Identification of 50 ALC-specific marker genes

We used the same approach to identify testicular marker genes specific to ALCs. Among the 120 genes specifically enriched in ALCs (compared to FLCs), 50 genes were considered as specific markers of ALCs (**Figure 5A-G** and **Table 2**). Among these 50 genes, we found *Hsd17b3* and *Hsd3b6*, two known markers of ALCs confirming our analysis. The remaining 48 genes are described for the first time as ALC specific markers. The analysis of the GO terms highlighted cellular processes such as protein transformation (*Klk1b21*, *Klk1b24*, *Klk1b27*, *C1rl*), peptidase regulation (*Serpina3c*, *Serpina3g*, *Serpina3n*, *Serpina5*) as well as metabolic processes (*Hsd17b3*, *Sult1e1*, *Bhmt*). To validate further our analysis, the specific expression of ALC marker genes *Bhmt* and *Sult1e1* was confirmed by *in situ* hybridisation (**Figure 5H-K**). We found that *Bhmt* and *Sult1e1* are co-expressed with *Star* in adult P100 testis but not in fetal E18.5 testis confirming that these genes are specifically expressed in ALCs and not in FLCs or any other testicular cells.

**Figure 5:**
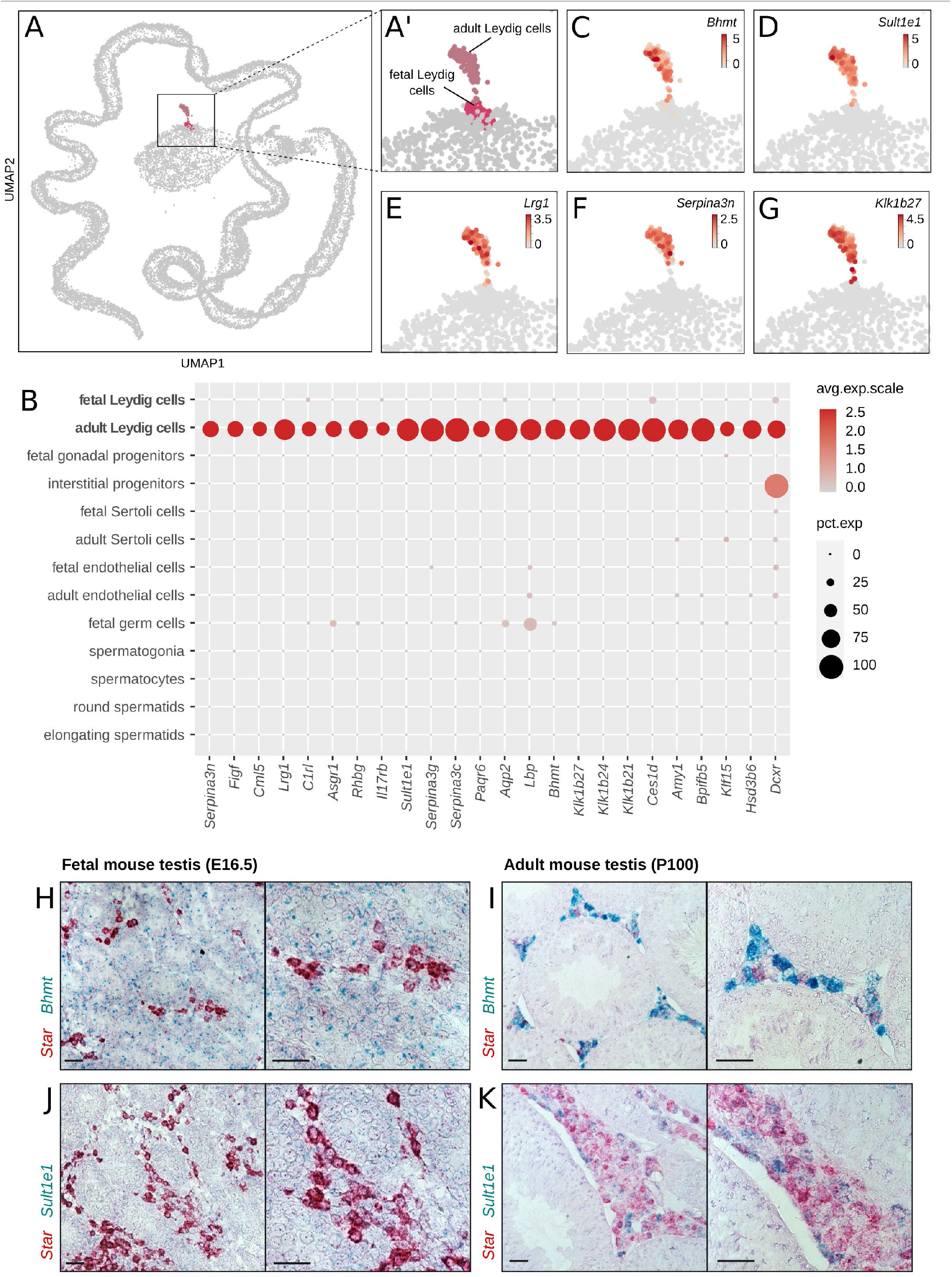
Identification of genes labelling ALCs. **(A, A’)** UMAP representation of single-cell transcriptomes highligting the two Leydig cell populations: fetal Leydig cells (FLCs) in pink and adult Leydig cells (ALCs) in pale violet. The other populations of the testis are colored in grey. **(C-G)** Enlargement on the UMAP representation colored according to the normalized expression of selected ALC-specific candidate marker genes, like *Bhmt* **(C)**, *Sult1e1* **(D)**, *Lrg1* **(E)**, *Serpina3n* **(F)**, and *Klk1b27* **(G)**. **(B)** Dotplot displaying the scaled expression of the top 25 ALC-specific candidate genes. **(H-K)** *In situ* hybridization assay by RNAscope^®^ labelling in red *Star*, a known marker of Leydig cells, and in green the ALC marker *Bhmt* **(H, I)** and *Sult1e1* **(J, K)** in adult mouse testis at P100 **(H, J)** and in embryonic mouse testis at E18.5 **(I, K)**. The colocalisation of the red and green probes indicate a co-expression of the two genes in the Leydig cells. The black scale bar corresponds to 100μm.

**Table 2.**
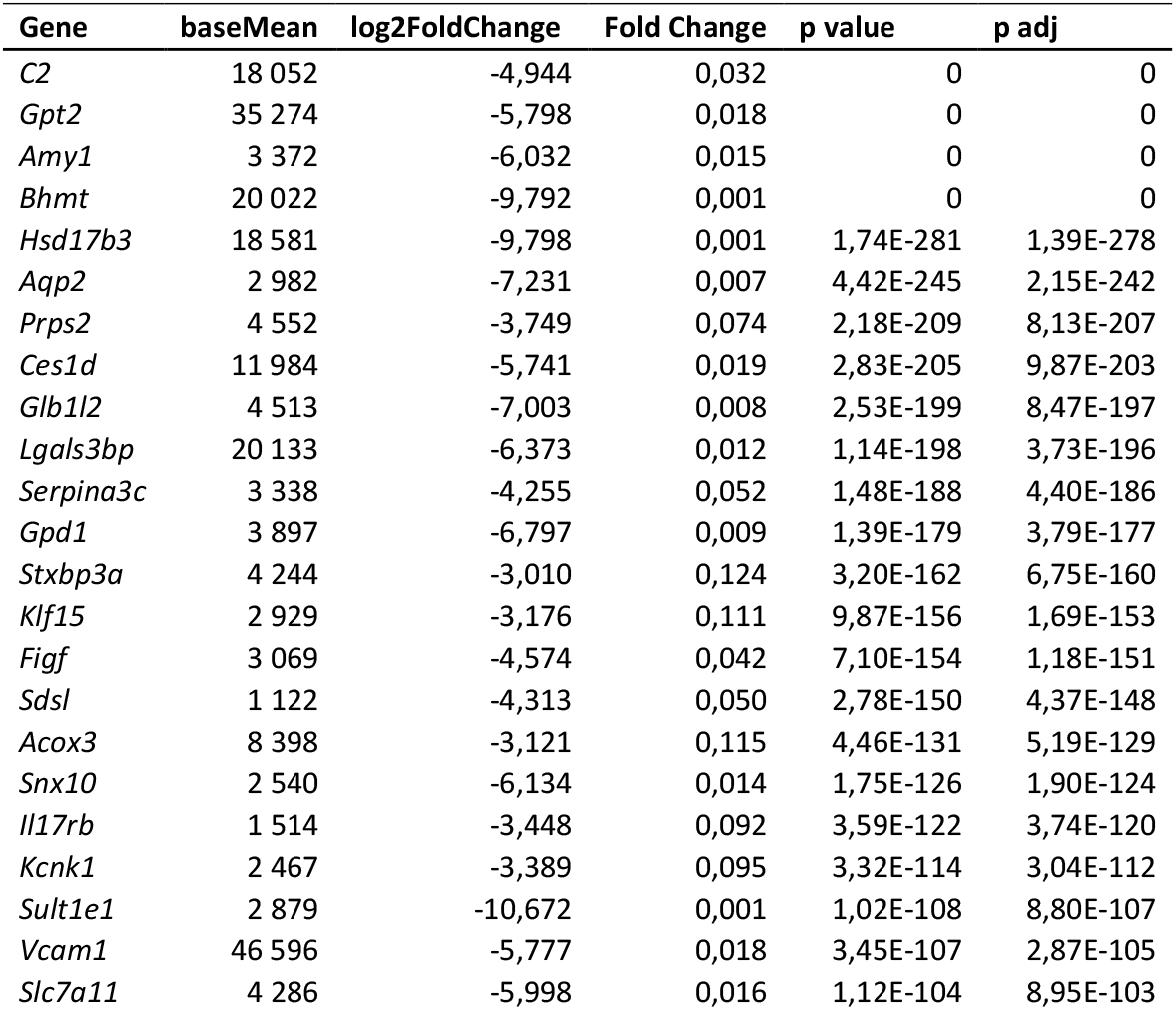
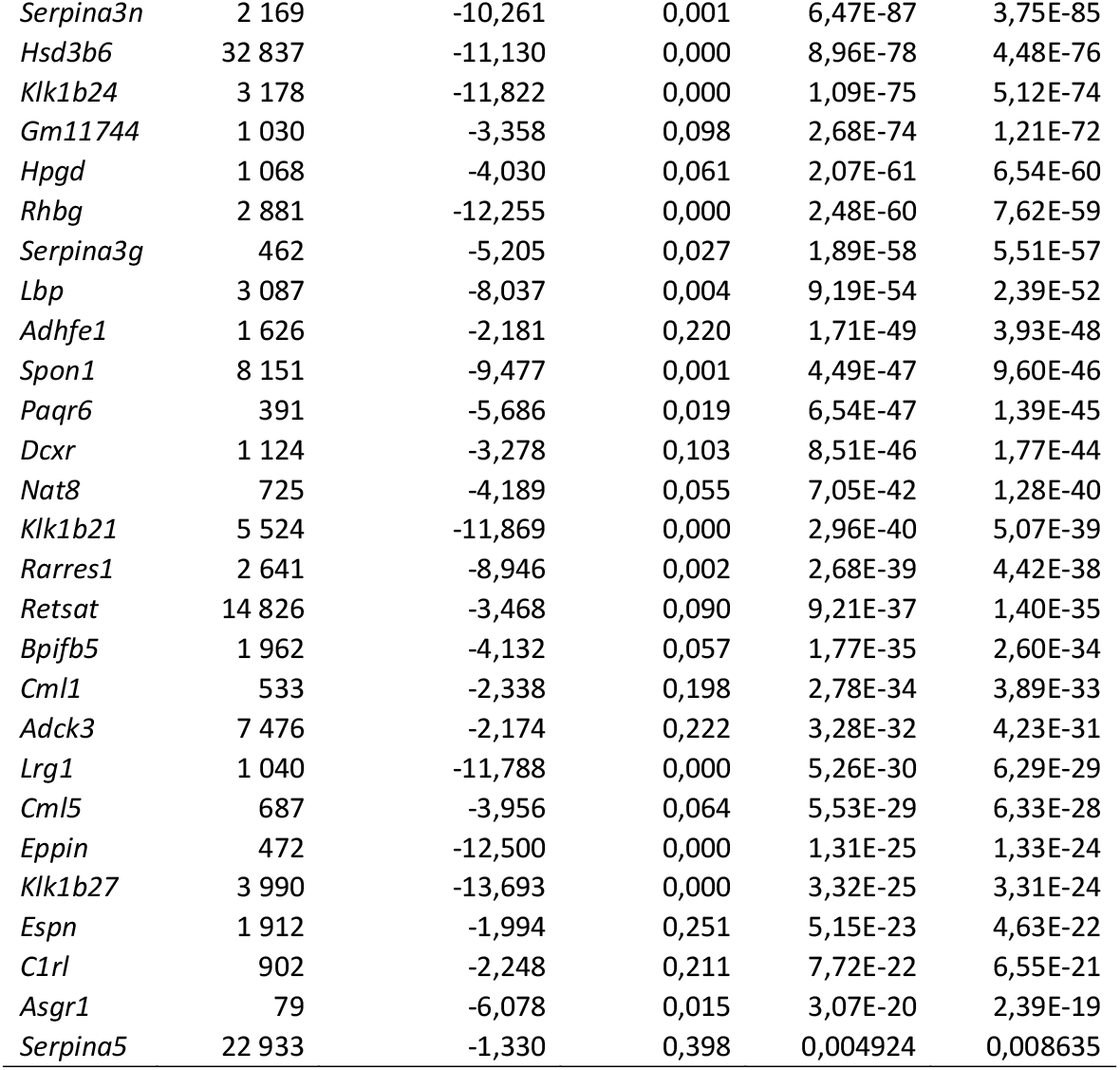
ALC specific genes

Overall, our analysis combining both bulk RNA sequencing and single-cell RNA sequencing resulted in the identification of 9 and 50 specific markers for FLCs and ALCs, respectively, most of which are newly identified.

## Discussion

The main purpose of our study was to characterize at transcriptomic level the similarities and differences between FLCs and ALCs, the major androgenic cells of the testis, using both bulk and single-cell RNA sequencing. Significant differences were observed both in terms of expression level, with 2,357 genes overexpressed in FLCs (11.2% of the total) and 3,396 genes overexpressed in ALCs (16.1%); and in terms of alternative splicing, with an over-representation of intron retention events in FLCs compared to ALCs. Our study also identified many specific markers for each Leydig cell populations, with 9 genes for FLCs and 50 genes for ALCs, most of them newly described.

### Identification of FLC- and ALC-specific genes

The purity of the Leydig cell population is critical for the identification of Leydig cell markers using microarray and bulk RNA sequencing analyses. We have multiple indications that support the assertion that the 9 FLC-specific markers - and 50 ALC-specific markers - identified in this study are robust and specific. First, our transcriptomic analysis combines two independent sources of data, namely those from the bulk RNA sequencing data of FLCs and ALCs, in which Leydig cells were sorted according to the level of GFP expression, but also a single-cell RNA sequencing of the testes of fetal and adult mice. In addition, the few markers already known, in particular *Hsd17b3* and *Hsd3b6* were also identified in the list of markers specific for ALCs. Finally, an independent validation by RNAscope^®^ of *Ren1* and *Crhr1* as FLC-specific marker genes, and *Bhmt* and *Sult1e1* as ALC-specific marker genes, confirmed their specific expression.

Several genome-wide expression studies using microarray technology or bulk RNA sequencing have investigated the transcriptome of FLCs (Jameson et al., 2012; McDowell et al., 2012; McClelland et al., 2015). In these studies, they isolated and evaluated the transcriptome of several populations present in the fetal testis, such as germ cells, Sertoli cells, Leydig cells and interstitial cells. Differential analysis of expression among these cell populations led to the identification of 166 overexpressed genes in FLCs. However, since the analysed cell populations represent only a fraction of the cell types present in the fetal testis, this list of FLC overexpressed genes is overestimated. In contrast, our analysis combining bulk and single cell RNA-seq identified nine FLC-specific candidates of which three were already described in these previous studies. We found that the other genes initially described as enriched in FLCs are mostly non-specific, either expressed in ALCs or in other testicular cell types. Among the three genes specifically enriched in FLCs are the *Crhr1* (McDowell et al., 2012), *Vsnl1* (Jameson et al., 2012; McClelland et al., 2015) and *Ren1* (Jameson et al., 2012) genes. The *corticotropin releasing hormone receptor 1* (*Crhr1*) is of particular interest as its ligand CRH (Corticotropin Releasing Hormone) is known to stimulate testosterone production in the fetal testes (McDowell et al., 2012) (see last paragraph of the discussion for more details and **Figure 6**). Moreover, our analysis revealed six new FLC-specific candidates including *Cyp26b1*, a gene coding for an enzyme degrading retinoic acid (RA), an active metabolite of retinol involved in meiosis regulation (Bowles et al., 2006; Koubova et al., 2006). It has recently been reported that in *Cyp26b1-/-* mutant mice, Leydig cell differentiation is impaired and steroidogenesis is decreased (Bowles et al., 2018).

**Figure 6:**
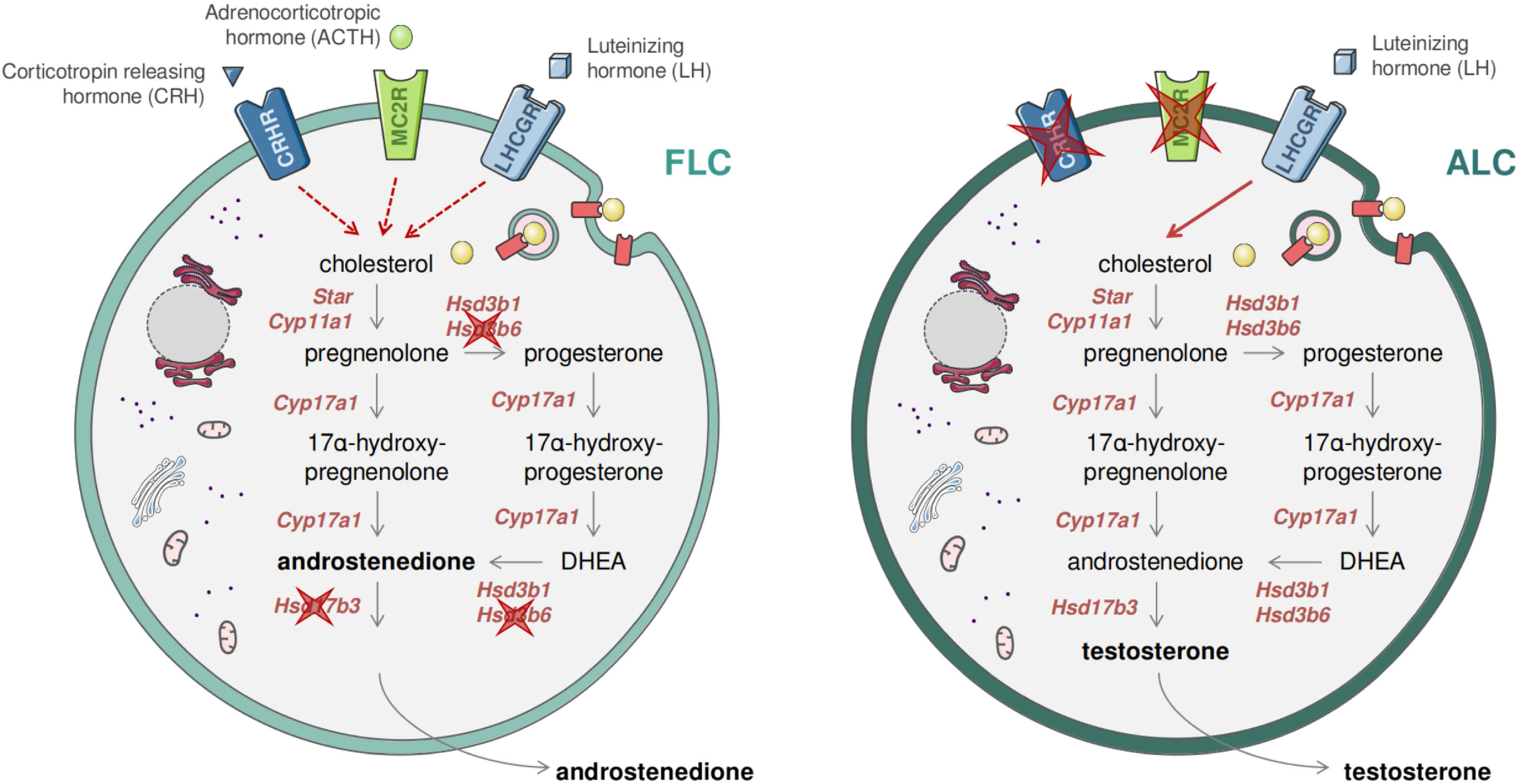
Dual and sequential model of steroidogenesis regulation. Drawing recapitulating the similarities and differences between FLCs (left panel) and ALCs (right panel). The expression of the ACTH receptor (encoded by *Mc2r*) and the CRH receptor (encoded by *Crhr1*) in FLCs suggests that the regulation of androgen production is mediated by LH, CRH and CRH, while in the absence of CRH and ACTH receptors in ALCs the regulation is exclusively under the control of LH.

Previous transcriptomics studies have used indirect methods to identify ALC-specific mRNA transcripts, such as the response of Leydig cells to hormones (Sanz et al., 2013), or cell ablation model using ethane dimethane sulphonate (EDS) to ablate LCs in adult male rats (O’Shaughnessy et al., 2014). These studies have resulted in a combined list of over 2000 genes whose expression is enriched in ALCs. Although ingenious, these approaches due to their technical bias and limitations do not guarantee an ALC-specific expression. Here, we have identified 50 genes with ALC-specific expression (**Table 2**). Confirming our results, three genes known to be specific for ALCs, *Hsd3b6*, *Hsd17b3* and *Vcam1* are also present in our list (O’ Shaughnessy et al., 2000; Shima et al., 2013; Wen et al., 2014). Among these 50 genes, six were not described in the previous studies, namely *Ces1d*, *Serpina3g*, *Rarres1*, *Bpifb5*, *Eppin*, and *Espn*. Hundreds of genes initially described as enriched in ALCs by previous studies were excluded because their expression was not specific. This is particularly the case for *Ptgds* and *Hsd11b1*, two genes often described as specific to ALCs (Baker and O’Shaughnessy, 2001; Wen et al., 2014). In this study, we validated by RNAscope^®^ the ALC-specific expression of two genes: *Betaine-Homocysteine S-Methyltransferase* (*Bhmt*) and *Sulfotransferase Family 1E Member 1* (*Sult1e1*). We also proved the specific expression of BHMT at the protein level by immunohistochemistry (**Supplementary Figure S2**). BHMT plays a key role in regulating betaine concentration, that can be stored to control cellular osmolarity or metabolised to provide a methyl group for homocysteine methylation (Alirezaei et al., 2012). It has been shown that the testes are among the organs that contains the most betaine (Slow et al., 2009). SULT1E1, for its part, plays a protective role for Leydig cells and seminiferous tubules against oestrogen overstimulation by catalysing the sulfo-conjugation and inactivation of oestrogens (Song, 2007). It is also noteworthy to find several members of the SERPIN family among our ALC-specific genes (*Serpina3c*, *Serpina3g*, *Serpina3n*, *Serpina5*) as most of them have been found in Leydig cells and seem to be sensitive to hCG (human chorionic gonadotropin) (Odet et al., 2006).

### Alternative splicing: high intron retention in FLCs

The main types of alternative splicing are alternative exon usage, alternative 5’ or 3’ splice sites, mutually exclusive exons, and intron retention. Intron retention is characterized by the inclusion of one or more introns in mature mRNA transcripts and has been previously considered to be an artefact of a dysfunctional spliceosome. It is known that alternative splicing, including intron retention, is frequent during embryonic development and contribute not only to the plasticity of the transcriptome but also the regulation of gene expression and protein diversity (Revil et al., 2010; Kalsotra and Cooper, 2011; Grabski et al., 2021). Here, we showed that intron retention is a landmark of the FLC transcriptome. Messenger RNA displaying intron retention are generally restricted from exiting the nucleus. This was proposed as a mechanism to downregulate gene expression (Grabski et al., 2021). In FLCs, we also showed that genes presenting alternative exon skipping are involved in splicing regulation itself. The control expression levels and activities of RNA binding proteins (RBPs) that regulate RNA splicing is mediated by auto-regulatory feedbacks by directly influencing the splicing of their own mRNAs (Müller-McNicoll et al., 2019). In particular, the regulation of the splicing factors of the SR (Serine/arginine rich) family regulate their activity by modulating the inclusion of a cassette exon containing a premature termination codon to produce or nor a functional protein (Müller-McNicoll et al., 2019). Tight regulation of the splicing factors is necessary for post-transcriptional gene expression regulation. The intron retention events observed in FLCs might subsequently result from the auto-regulation feedback of the splicing factors. Post-transcriptional gene expression regulation through alternative splicing have been identified as key player in the differentiation of mesenchymal stem cells (Park et al., 2020). We can postulate that the regulation of the FLC differentiation might also be mediated by alternative splicing.

### Differences in FLCs and ALCs transcriptomes affect steroidogenesis and its regulation

FLCs and ALCs display significant differences in both steroidogenic regulation and the type of androgen produced (androstenedione vs testosterone) (O’Shaughnessy et al., 2000, 2002; Shima et al., 2013). Our transcriptomic data confirmed the differences in androgen production, the expression of the *Hsd17b3* gene encoding the enzyme responsible for the conversion of androstenedione to testosterone is not expressed in FLCs but only in ALCs (**Table 2** and **Figure 6**), which explains why FLCs synthesise mainly androstenedione and ALCs are capable of producing testosterone (O’Shaughnessy et al., 2000; Rebourcet et al., 2020). Regarding the differences in steroidogenesis regulation, LH appears not to be essential for FLC function since androgen production and masculinization of the fetus occurs normally in LH/CG receptor knockout mice (Kendall et al., 1995; Lei et al., 2001; Zhang et al., 2001; Ma et al., 2004; O’Shaughnessy and Fowler, 2011; Teerds and Huhtaniemi, 2015). However in *T/ebp/Nkx2.1* null mice, which lack a pituitary gland, testicular androgen levels are markedly reduced in late gestation, suggesting that additional hypothalamo/pituitary hormones may be required for Leydig cell function and androgen production. Interestingly, our transcriptomic analysis revealed that the ACTH receptor, melanocortin type 2 receptor (*Mc2r*), and the corticotropin releasing hormone receptor 1 (*Crhr1*) are both specifically expressed in FLCs and absent from ALCs (**Table 1** and **Figure 6**). ACTH has been reported to stimulate *in vitro* testosterone production in fetal but not in adult testes suggesting that FLCs, but not ALCs, are sensitive to ACTH stimulation(O’Shaughnessy et al., 2003). However, fetal testosterone levels were normal in *Proopiomelanocortin* (*POMC*)-deficient mice that lack circulating ACTH, indicating that ACTH, like LH, is not essential for FLC function. Corticotropin-releasing hormone (CRH) has been also reported to stimulate steroidogenesis by direct activation of FLCs in fetal rat and mouse testes *ex vivo* and in MA-10 mouse Leydig cells (McDowell et al., 2012). In contrast, CRH does not enhance steroidogenesis in primary ALCs (Huang et al., 1995; McDowell et al., 2012). Combined together, these results suggest a sequential regulation of steroidogenesis in LCs. In this model, androgen production by FLCs is stimulated by three potentially redundant hypothalamo/pituitary hormones, namely LH, CRH and ACTH. Fetal androgen production can occur in the absence of any of these hormones with the two other hormones able to maintain FLC steroidogenic activity. Conversely, activation of steroidogenesis in ALCs is LH-dependent and CRH- and ACTH-independent, since *Crhr1* and *Mc2r* are not expressed in these cells (**Figure 6**). Although this model of steroid regulation by Leydig cells needs to be confirmed by further studies, such a mechanism may have evolved to ensure the production of adequate levels of androgens during fetal development.

## Disclosure statement

The authors declare no conflicts of interest.

## Funding sources

This work was supported by the Swiss National Science Foundation (Grant 31003A_152636 to S.N.); and the Département de l’Instruction Publique of the State of Geneva (to S.N.). Department of Health | National Health and Medical Research Council (NHMRC), Grant/Award Number: APP1158344 (to L.S. & D.R.).

## Acknowledgments

The authors thank Violaine Regard from the NEF laboratory, Cécile Gameiro and Gregory Schneiter from the Flow Cytometry Facility, the Genomics Platform of iGE3, and the Animal Facility of the Faculty of Medicine of the University of Geneva.

---

**Supplementary Figure 1:**
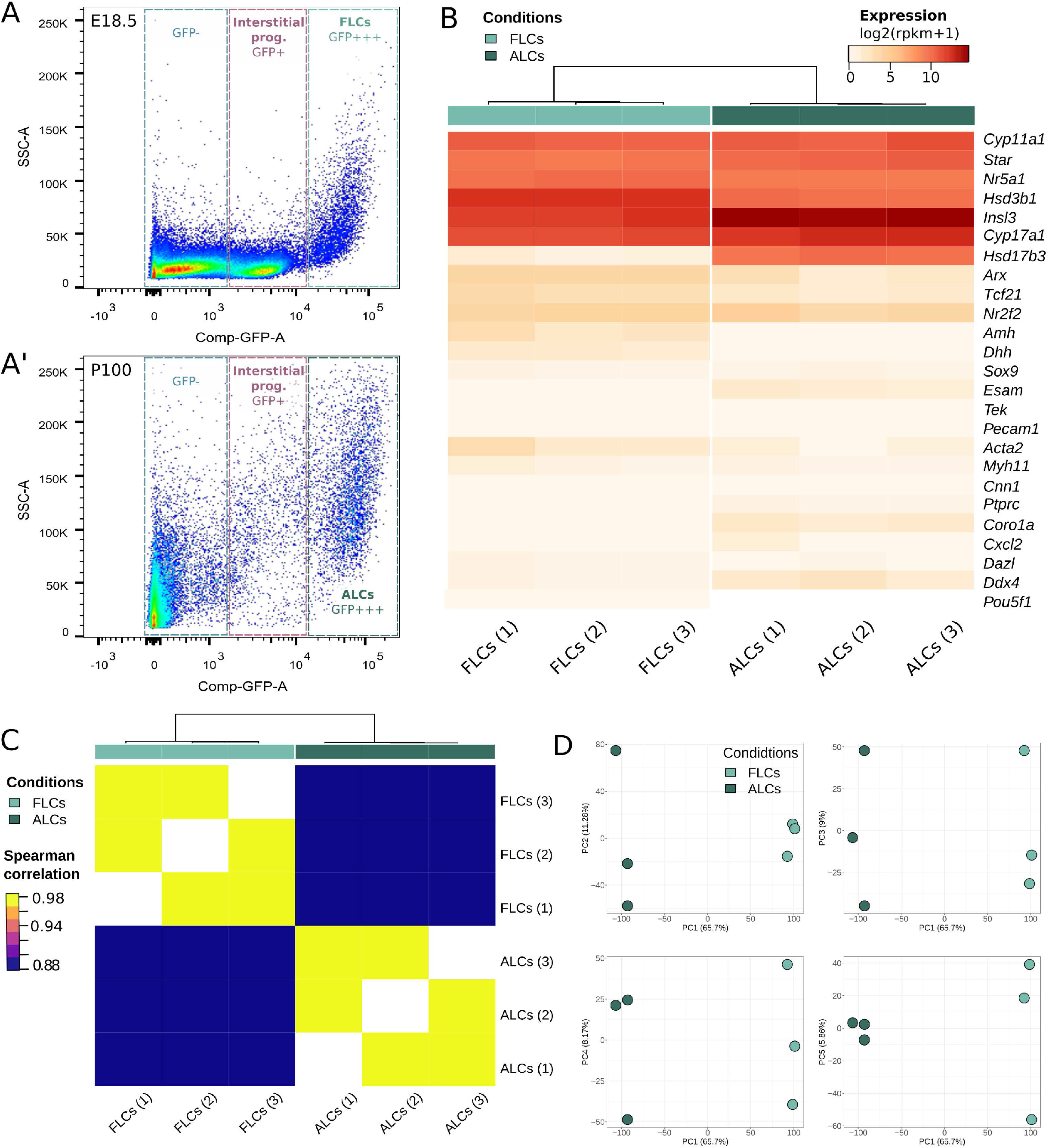
**(A,A’)** Scatter plot showing the selection of Leydig cells (GFP+++) by FACS in E18.5 **(A)** and in P100 **(A’)** testis cell suspension. The X axis corresponds to the GFP florescence level and the Y axis forward scatter area explaining the granularity of the events. **(B)** Heatmap showing the normalized scaled expression of selected marker genes in our three replicates at E18.5 (pale green) and in our three replicates at P100 (dark green). The color is representative of the expression level. The gene expression confirmed the purity of the samples in Leydig cells. *Cyp11a1*, *Star*, *Nr5a1*, *Hsd3b1*, *Insl3*, *Cyp17a1*, *Hsd17b3*, *eGFP*: Leydig cells. *Cdh5*, *Pecam1*, *Acta2*, *Rgs5*: Endothelial cells. *Pdgfra*, *Tcf21*, *Wnt5a*, *Nr2f2*: Interstitial progenitors. *Amh*, *Lhx9*, *Dhh*, *Sox9*: Sertoli cells. *Pou5f1, Mael, Dadx4, Dazl:* Germ cells. **(C)** Heatmap showing the Spearman correlation score between the six Leydig cells samples. The score is indicated by the color scale. **(D)** Principal Component Analysis (PCA) plot where each dot corresponds to a Leydig cells sample. The dots are colored according to the sample type. E18.5: pale green; P100: dark green.

**Supplementary Figure 2:**
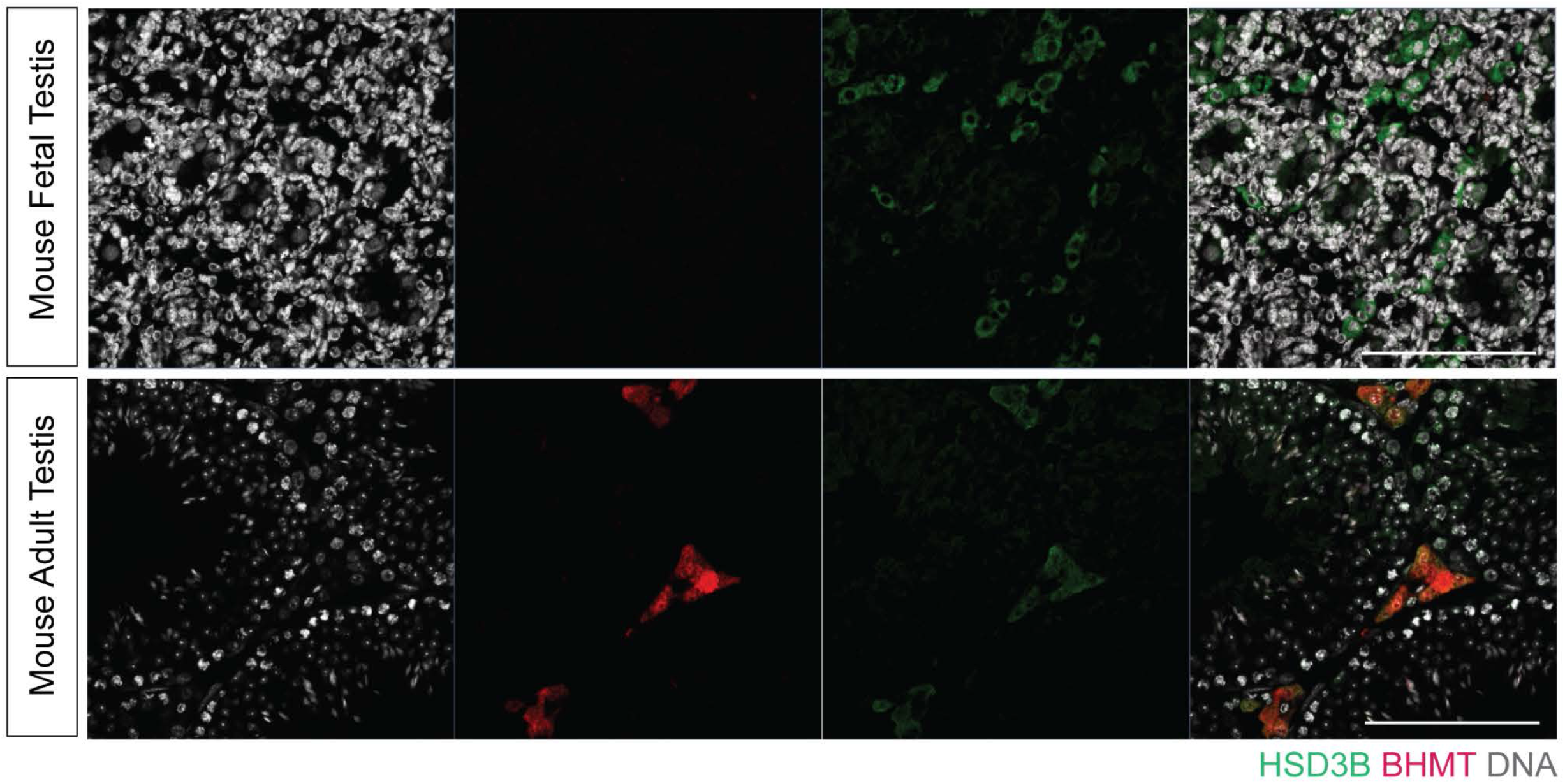
Immuno-histochemistry (IHC) staining on mouse fetal and adult testis. The DNA is colored in white, HSD3B labels Leydig cells in green and BHMT is colored in red. The white scale bar corresponds to 50μm.

## Notes

### Competing Interest Statement

The authors have declared no competing interest.

